# Seizure localisation with attention-based graph neural networks

**DOI:** 10.1101/2020.12.03.409979

**Authors:** Daniele Grattarola, Lorenzo Livi, Cesare Alippi, Richard Wennberg, Taufik A. Valiante

**Affiliations:** Faculty of Informatics, Università della Svizzera italiana, Lugano, Switzerland; Departments of Computer Science and Mathematics, University of Manitoba, Winnipeg, Canada; Department of Computer Science, University of Exeter, Exeter, United Kingdom; Department of Electronics, Information, and Bioengineering, Politecnico di Milano, Milan, Italy; Division of Neurology, Department of Medicine, Krembil Brain Institute, Toronto Western Hospital, University of Toronto, Toronto, Canada; Department of Surgery, Division of Neurosurgery, University of Toronto, Toronto, Canada; Krembil Brain Institute, Division of Clinical and Computational Neuroscience, Toronto, Canada; Institute of Medical Sciences, University of Toronto, Toronto, Canada; Institute of Biomedical Engineering, University of Toronto, Toronto, Canada; Electrical and Computer Engineering, University of Toronto, Toronto, Canada

## Abstract

In this paper, we introduce a machine learning methodology for localising the seizure onset zone in subjects with epilepsy. We represent brain states as functional networks obtained from intracranial electroencephalography recordings, using correlation and the phase locking value to quantify the coupling between different brain areas.

Our method is based on graph neural networks (GNNs) and the attention mechanism, two of the most significant advances in artificial intelligence in recent years. Specifically, we train a GNN to distinguish between functional networks associated with interictal and ictal phases. The GNN is equipped with an attention-based layer that automatically learns to identify those regions of the brain (associated with individual electrodes) that are most important for a correct classification. The localisation of these regions does not require any prior information regarding the seizure onset zone.

We show that the regions of interest identified by the GNN strongly correlate with the localisation of the seizure onset zone reported by electroencephalographers. We report results both for human patients and for simulators of brain activity. We also show that our GNN exhibits uncertainty on those patients for which the clinical localisation was unsuccessful, highlighting the robustness of the proposed approach.

## I. Introduction

Epilepsy is a neurological disorder characterised by recurrent episodes of excessive neuronal firing (Stafstrom and Carmant, 2015). In approximately a third of the patients, epilepsy cannot be treated with anti-seizure drugs and resective surgery can be considered as a possible treatment (Kwan and Brodie, 2000). The outcome of surgery is crucially dependent on the successful localisation of the seizure onset zone (SOZ) (Burns et al., 2014; Van Mierlo et al., 2014).

Electroencephalography (EEG) is the mainstay for studying and diagnosing epilepsy, and it is widely used to detect, classify, and localise seizures by recording and processing the electrical activity of groups of neurons (Nunez et al., 2006). However, due to their low spatial resolution, scalp EEG recordings in some cases are not informative enough to successfully localise seizures (Shah and Mittal, 2014). In these cases, intracranial EEG recordings (iEEG), in which electrodes are placed directly on or within the brain, provide better spatio-temporal resolution to capture the dynamics of seizure generation and propagation (Hashiguchi et al., 2007). However, the high temporal resolution of iEEG and the complex functional interaction of distant brain areas, especially during seizures, make the interpretation and processing of raw iEEG data a non-trivial task for clinicians. For this reason, a significant branch of epilepsy research is concerned with summarising iEEG data by considering the pairwise (statistical) dependencies between the activity of different brain areas over time (Van Mierlo et al., 2014). These dependencies are usually represented by *functional networks* (FNs), in which each node represents a sensor and edges are weighted by a *functional connectivity* (FC) metric (Bastos and Schoffelen, 2016).

FNs are a widespread tool to study seizure localisation, with early approaches dating back to the 1970s (Gersch and Goddard, 1970; Brazier, 1972). Seizures have been observed to affect the functional organisation of brain activity at the mesoscale, both from a node-centric (Burns et al., 2014) and an edge-centric (Khambhati et al., 2015) perspective. In particular, Burns et al. (2014) identified sets of brain states that emerge by clustering FNs, consistent in interictal and ictal periods for individual patients. They observed that changes in node centrality in FNs accurately predict the SOZ. Khambhati et al. (2015) observed a strengthening of FC in the SOZ during seizures, also coinciding with a topological tightening of the connections (*i.e.*, strong connections also become physically closer). Khambhati et al. (2016) proposed *virtual cortical resection*, *i.e.*, the removal of nodes from FNs, in order to study changes in network synchronizability, which is a known predictor for the spread of seizures (Schindler et al., 2008). Lopes et al. (2017) also observed that the resection of brain areas associated with *rich-club* hubs in FNs correlates with a good postoperative outcome. Seizure localisation has also been studied in FNs obtained from functional magnetic resonance imaging (fMRI) (Lee et al., 2014; Weaver et al., 2013) and scalp EEG (Staljanssens et al., 2017) data. Recent work by Covert et al. (2019) used spatio-temporal graph convolutional networks (ST-GCNs) (Yu et al., 2017) to perform seizure detection. They conducted an *expost* analysis similar to that of Khambhati et al. (2016) to quantify the importance of a node by observing the effect of its removal on the downstream detection accuracy. Gadgil et al. (2020) also proposed a methodology based on ST-GCNs to identify high-interest areas in fMRI by learning to estimate edge importance, although they did not apply it to seizure localisation. For a more in-depth review of approaches to seizure localisation with FNs, we refer the reader to Van Mierlo et al. (2014).

This paper aims to use the representation of brain states as FNs to automate the localisation of seizures using deep learning. Advances in deep learning techniques over the past decade have revolutionised how high-dimensional, high-volume data can be used in the context of artificially intelligent systems. In particular, deep learning techniques for computer vision have shown how artificial intelligence can be successfully adopted in clinical settings to aid human experts in their decision making (Litjens et al., 2017). Despite these successes, traditional deep learning methods are limited to processing regular structures like images and time series, and cannot naturally consider the relations that exist in a complex system with multiple interacting components, such as those described by FNs evolving over time. For this reason, recent literature has seen the rise of Graph Neural Networks (GNNs) (Battaglia et al., 2018; Bronstein et al., 2017) as a generalisation of deep learning techniques to process data represented as arbitrary graphs.

In this paper, we introduce a GNN-based methodology for seizure localisation, using FNs to efficiently represent brain states. The core of our algorithm is a GNN equipped with an *attention-based readout*. By training the GNN to perform seizure detection, the readout automatically learns to pay more attention to those nodes that are more important for a correct classification. Then, we propose a simple and fast way of analysing the attention coefficients over time, so that we obtain a ranking of the nodes based on their overall importance in detecting a seizure. Crucially, our methodology does not require *a priori* information regarding the SOZ, but only weak supervision in the form of annotated seizure onsets and offsets. A schematic representation of our approach is shown in Figure 1.

**Fig. 1:**
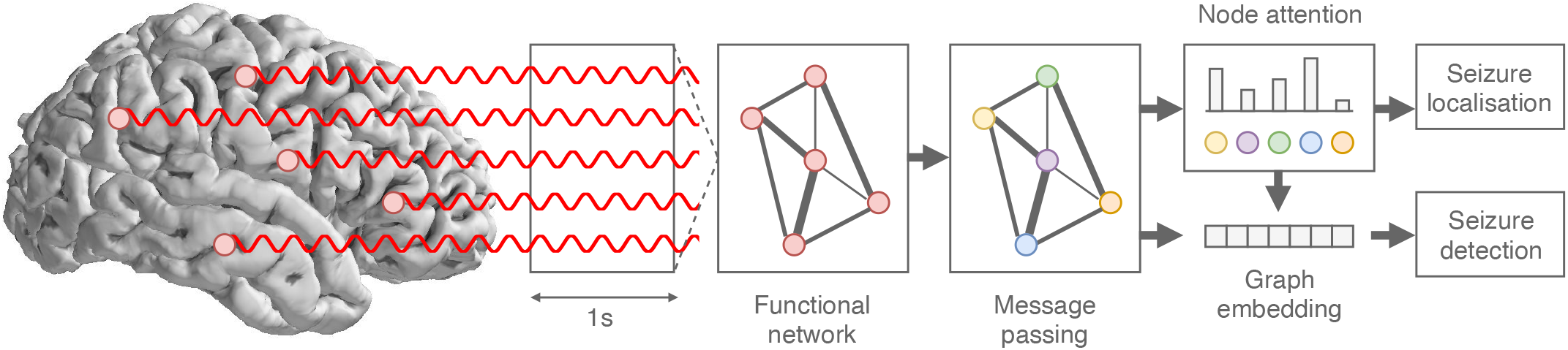
Schematic view of our GNN-based pipeline for seizure detection and localisation. Starting from raw iEEG data, we compute a functional network to represent the spatio-temporal dynamics of the signals compactly. The FN is then given as input to a GNN composed of an edge-aware message passing operation followed by an attention-based readout to compute a graph-level embedding. The embedding is then classified to perform seizure detection, while the attention scores are analysed to perform seizure localisation.

We validate the proposed methodology on clinical iEEG data collected from eight human subjects and show that the electrode rankings computed with our localisation procedure are highly correlated with the true SOZs. We also validate our algorithm on simulated data, using a simple model of seizure initiation (Benjamin et al., 2012) and a more complex brain simulator (Sanz Leon et al., 2013) based on the Epileptor model (Jirsa et al., 2014). Our main contributions and results are summarised as follows:

- We present a new algorithm for seizure localisation based on GNNs, which uses FNs to represent brain states in a compact form and requires no explicit supervision on the SOZ;
- We show that the attention coefficients learned by the GNN correlate with clinically-identified SOZs and accurately predict the presence of ictal activity;
- We show that, when electroencephalographers were not able to identify the SOZ from the iEEG data, the GNN also shows uncertainty in the localisation;
- We show that, as expected, the choice of FC metric used to estimate FNs is important for an accurate localisation;
- Finally, we show that our methodology performs well on very imbalanced datasets, achieving a good localisation accuracy even on patients for which we observe as few as five seizures during training.

## II. Methods

### Notation

We denote a time series *x_i_*(*t*) to represent the *i*-th iEEG channel at time *t*. We define a graph as a tuple 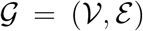, where 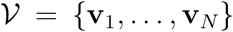 represents the set of attributed nodes with attributes **v**_*i*_ ∈ ℝ^*F*^, and 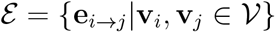 represents the set of attributed edges with attributes **e**_*i→j*_ ∈ ℝ^*S*^ indicating a directed edge between the *i*-th and the *j*-th node. We indicate the neighbourhood of node *i* with 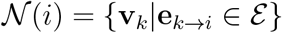. We say that a graph is undirected if **e**_*i*→*j*_ ∈ *ε* ⇔ **e**_*j*→*i*_ ∈ *ε*. Note that in the text, for simplicity, we refer to nodes using their index, *e.g.*, node *i*.

#### A. Functional networks

Choosing a suitable FC metric to model the pairwise interaction between brain areas is a non-trivial challenge, as there exist a large variety of methods with their advantages and disadvantages. FC metrics can be characterised according to several properties, including whether they are in the time or frequency domain, whether they are directed or undirected (*i.e.*, if they model asymmetric or symmetric couplings), or whether they are model-free or model-based (Bastos and Schoffelen, 2016). Here, we focus on undirected FC metrics to simplify the GNN computation, and on model-based approaches to reduce the computational costs of estimating the FC metrics directly from data. We do, however, consider two different metrics to highlight the practical differences that emerge between time- and frequency-domain metrics.

FNs are generated by computing a FC value for each pair of iEEG channels *x_a_*(*t*) and *x_b_*(*t*) over a time window of length *T*. For the time-domain metric, we consider Pearson’s correlation coefficient:

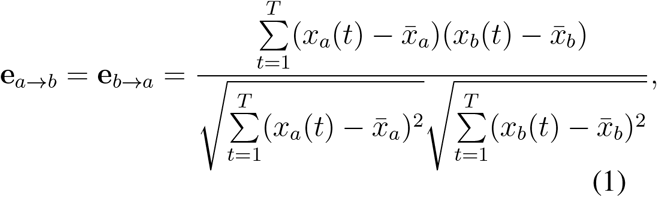

where 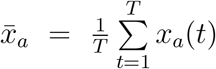 and analogously for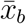. Correlation allows to quantify symmetric linear interactions, it is easy to compute and, as such, it is often used in the literature. For the frequency domain, we consider the phase-locking value (PLV) (Lachaux et al., 1999):

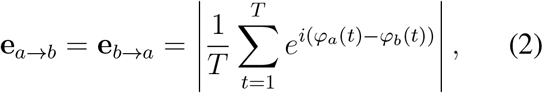

where *φ_a_*(*t*) indicates the instantaneous phase of signal *x_a_*(*t*) obtained via Hilbert transform (and similarly for *φ_b_*(*t*)). A significant advantage of PLV over correlation is that it is less sensitive to artefacts in the iEEG signals (such as those caused by the patient’s movements). After computing the FC metrics for each pair of channels, we sparsify the resulting FNs by removing those edges for which |**e**_*i*→*j*_| < 0.1, *i.e.*, those indicating weak coupling. The choice of sparsification threshold is generally an important hyperparameter when studying FNs. For example, a principled way of computing a dynamic sparsification for each individual FN is described in the work of Kramer et al. (2009). However, in this case, we are not interested in fine-tuning the threshold nor do we wish to devise a dynamic sparsification scheme to process each FN independently. As long as the same threshold is consistently used for different FNs, then the GNN will learn to deal with the resulting distribution of FNs. We report an additional discussion regarding the threshold in the Appendix.

We generate a dataset of FNs for each patient, dividing the FNs into ictal and interictal classes and proceeding in a per-seizure fashion. Let *f_s_* be the sampling rate of the iEEG signal, *L* the duration of a seizure, *t*_0_ the time indicating the seizure onset, *k* ≥ 1 a subsampling factor, and *T* the length of the time windows. Additionally, let *y*(*t*) ∈ {0, 1} be a binary signal indicating whether the patient is having a seizure at time *t* (*i.e.*, *y*(*t*) = 1 if *t ≥ t*_0_ and 0 otherwise). Note that we consider each seizure to end at time *t*_0_ + *L* and we do not compute FNs for the data immediately following a seizure offset.

Given a time window [*t* − *T,…, t*], we compute a FN 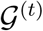 and label it with class

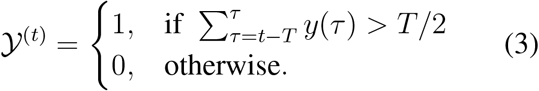

To generate the FNs associated with seizures (class 1), we consider the data interval [*t*_0_ − *T/*2*, …, t*_0_ + *L*] and take overlapping windows of size *T* with a stride of 1*/f_s_*. For the interictal FNs (class 0), instead, we consider a longer period preceding the seizure onset, [*t*_0_ − *kL, …, t*_0_ + *T/*2], and we take windows at a larger stride of *k/f_s_*. In this work, we consider *k* = 10 and *T* = 1s for all experiments, although other values are possible.

This procedure to generate the FNs (summarised in Figure 2) results in a balanced dataset and has two advantages. First, it allows us to fully use all the available (and rare) ictal events. Second, it allows us to consider a more diverse sample for the interictal class. The small differences between consecutive FNs of the positive class, due to the small stride at which windows are taken, can be seen as a form of sample weighting to account for the class unbalance characterising the problem.

**Fig. 2:**
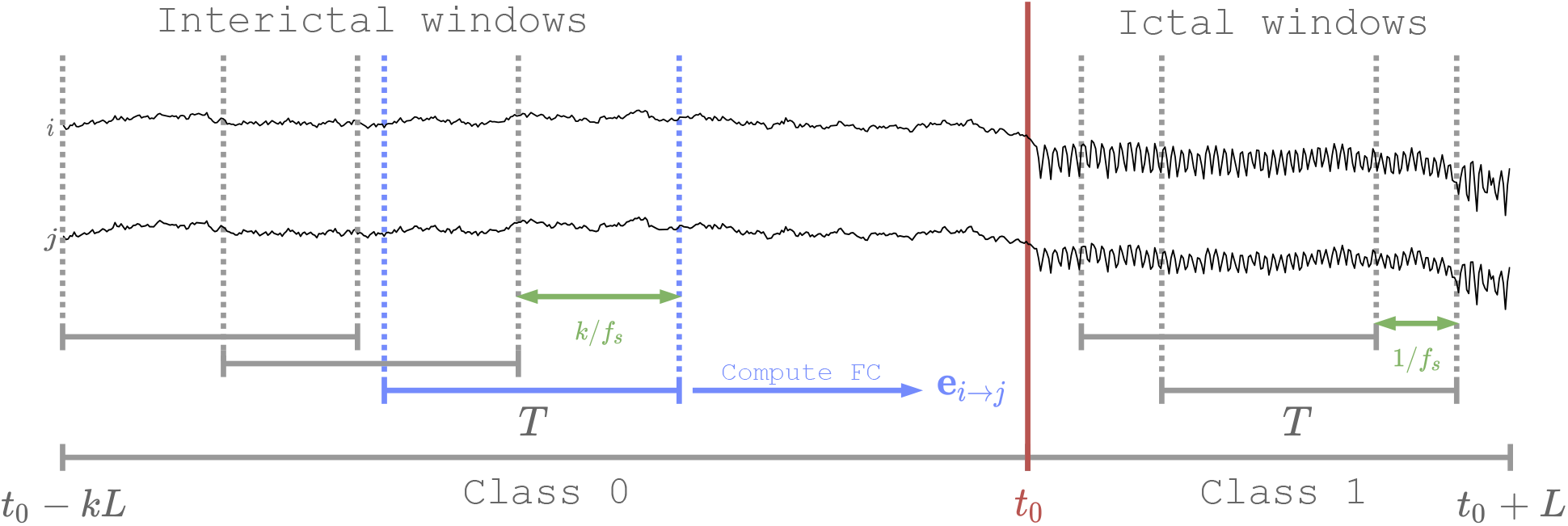
Schematic representation of the procedure used to generate FNs. For each seizure of length *L* starting at *t*_0_ (marked in red), we consider an interictal interval of length *kL*. Interictal FNs are generated taking windows of length *T* at stride *k/f_s_*, while ictal windows are taken with stride 1*/f_s_* (in green). For each window and each pair of electrodes *i* and *j*, we compute the FC value **e**_*i→j*_ (in blue) to obtain the full FN. This figure is only meant to represent the procedure and is not shown in any physical temporal scale.

In order to have initial node features that can be processed by the GNN, we consider dummy attributes set to 1 for all nodes. Other choices that depend on the actual iEEG signals are possible (*e.g.*, the signal power or wavelet coefficients) but were not explored in this work.

#### B. Attention mechanism

Attention (Bahdanau et al., 2014; Vaswani et al., 2017) is a processing technique for neural networks to learn how to selectively focus on parts of the input. Originally developed for aligning sentences in neural machine translation (Bahdanau et al., 2014; Vaswani et al., 2017), the attention mechanism has been used to achieve state-of-the-art results on different tasks like language modelling (Brown et al., 2020), image processing (Xu et al., 2015), and even learning on graphs (Velickovic et al., 2018).

In this paper, we focus on the concept of *self* - attention, which indicates a class of attention mechanisms that learn to attend to the output of a layer using the output itself (in contrast to classical attention, which uses the output of one layer to focus on the output of another – *e.g.*, the sentence of the source language is used to focus on the target language). At its core, self-attention consists of computing a compatibility score *α_ij_* ∈ [0, 1] between two vectors **h**_*i*_, **h**_*j*_ ∈ ℝ^*F*^ (both part of the same sequence, image, graph, etc.):

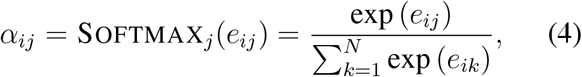

where

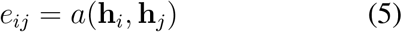

and *a* is called an *alignment* model, which is usually learned end-to-end along with the other parameters of the neural network. The compatibility score is then used to compute a representation of element *i* as:

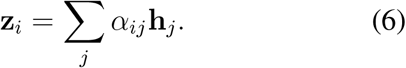

Intuitively, the attention mechanism learns the importance of element *j* to describe element *i*, and computes score *α_ij_* to quantify this importance. The alignment model can be seen as a similarity function between the two elements, which is then normalised via the Softmax function. Different implementations of the alignment model are possible, although often it is implemented as a multi-layer perceptron.

Attention mechanisms are usually trained without direct supervision and automatically learn to focus on different parts of the data according to the loss of the given task. By optimising the overall task loss, the attention layers in a neural network learn to compute the optimal compatibility scores. This is a key aspect of our proposed methodology, where we use self-attention to automatically detect those brain areas (monitored via different iEEG channels) that are important to detect a seizure. We stress that, crucially, using attention allows us to perform localisation without ever providing our neural network with ground truth information on the SOZ.

#### C. Graph neural networks for seizure localisation

Graph Neural Networks (GNNs) are a class of neural networks designed to perform inference on graph-structured data (Battaglia et al., 2018). At their core, GNNs learn to represent the nodes of a graph by propagating information between connected neighbours, whereas a global representation of the entire graph is usually obtained by computing a *readout* of the nodes, like a sum, average, or component-wise maximum vector. In this work, we focus on the family of *message-passing* networks (Gilmer et al., 2017), in which the *l*-th layer maps the attributes 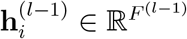 of the *i*-th node to:

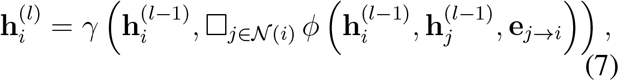

where 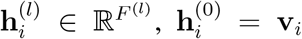, and *ϕ* and *γ* are differentiable functions equivariant to node permutations, respectively called the *message* and *update* functions, while □ is a permutation-invariant function (such as the sum or the average) to aggregate incoming messages.

Many recent papers have introduced methods for graph representation learning based on this general scheme, with different implementations ranging from polynomial (Defferrard et al., 2016) or rational (Bianchi et al., 2019) graph convolutional filters, to attentional mechanisms (Velickovic et al., 2018). In most of these works the creation of messages is only dependent on the node attributes, although some methods have been proposed that also explicitly take edge attributes into account (Simonovsky and Komodakis, 2017; Schlichtkrull et al., 2018). In particular, the Edge-Conditioned Convolutional (ECC) operator proposed by Simonovsky and Komodakis (Simonovsky and Komodakis, 2017) incorporates edge attributes into the message-passing scheme by using a *kernel-generating network f* ^(*l*)^(·) that dynamically computes messages between each pair of connected nodes. An ECC layer is thus defined as:

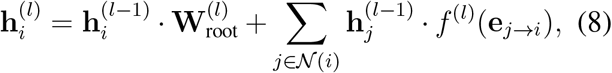

where 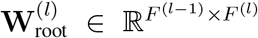 is a learnable kernel applied to the root node itself and the kernel-generating network is usually a multi-layer perceptron 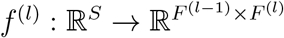.

Our method for seizure localisation can be summarised as follows. First, we train a GNN with an attention-based readout to detect seizures from FNs. This is a graph-level classification problem where a label (ictal or interictal) is assigned to each FN. Then, we analyse the compatibility scores learned by the attentional mechanism to identify those nodes that the model consistently considers as important. Although we train the GNN to do seizure detection in a supervised way, *i.e.*, it requires manually-annotated seizure onsets and offsets, the localisation is fully unsupervised. This is one of the main strengths of the proposed method, as significantly less manual work is required to annotate the temporal boundary for each seizure, rather than the SOZ.

There are two main components in our GNN architecture. First, the connectivity information is propagated to the node attributes via an edge-aware message-passing operation like ECC. A single layer is sufficient because the input FNs are densely connected, and most nodes will receive information from the whole graph in a single step of message passing.

Then, we use a self-attentional mechanism to compute the graph readout:

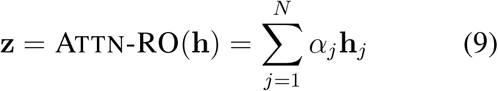

where

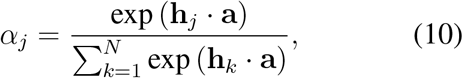

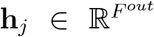 is the embedding of the *j*-th node computed by the ECC layer, and 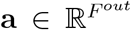 is a vector of learnable weights. Note that, compared to Equation (6), here index *i* is left implicit as the attention is only computed once for all nodes, to reduce the graph to a vector. This is also reflected in the fact that the alignment model is a function of only one node at a time, *e.g.*, **h**_*j*_ · **a**. For a more general way of applying attention to every possible pair of nodes (while maintaining the graph structure), see (Velickovic et al., 2018).

Finally, a multi-layer perceptron MLP(·) with sigmoid activation computes the probability that the input FN represents an ictal window of iEEG data.

The full architecture is written as:

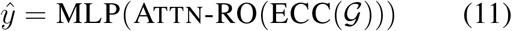

where 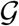 represents an input FN (*cf.* Figure 1).

By training the GNN to correctly distinguish the ictal FNs from the non-ictal ones, we also implicitly train the attentional readout Attn-RO to assign higher attention to those nodes of the FNs that maximise the confidence in the prediction. We then analyse how the attention scores assigned to nodes change over time, and rank the nodes according to the overall amount of attention that they receive before and during a seizure. The localisation procedure is described in the following section.

#### D. Localising the seizure onset zone

For each seizure in the data, we consider symmetric intervals of length 2*L* centred at the seizure onset, so that the first *L* timesteps are pre-ictal and the remaining *L* cover the beginning of the seizure. For each of the 2*L* timesteps, we compute a 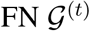 from a *T* = 1s window ending at time *t*, obtaining a sequence of FNs 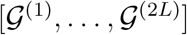 (this is equivalent to how we generate the training datasets, except that the subsampling is set at *k* = 1). For each FN in the sequence, we use the GNN to compute the attention scores over the nodes according to Equation (10). We thus compute a sequence of attention scores 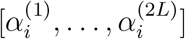 for each node *i*.

We then sum the sequence of attention scores to obtain the overall *importance* of the node over the considered time interval:

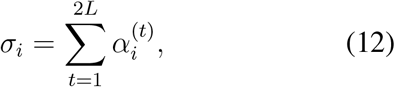

and normalise the importance scores to the [0, 1] interval as:

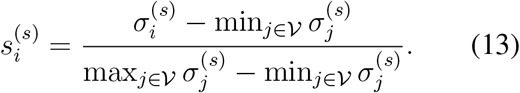

Finally, we rank the nodes according to their importance and predict the SOZ accordingly.

## III. Results

We report the results obtained on real iEEG data collected from eight patients. Additional results on two brain activity simulators (a simple network model (Benjamin et al., 2012) and *The Virtual Brain* simulator (Sanz Leon et al., 2013)) and all experimental details regarding the GNN are reported in the appendix.

### A. Data collection and pre-processing

We used iEEG data recorded from eight human subjects with medically refractory epilepsy, the recordings obtained as part of their standard clinical pre-surgical investigations. The patients were selected among a larger pool of patients based on certain criteria, chiefly having at least 5 clinical seizures recorded in our database and having a recorded clinical history of at least 2 years.

The study was approved by the Research Ethics Board at the University Health Network (ID number 12-0413) and written consent for data collection was obtained from all participants. Each patient had a varying number of recorded clinical seizures and the number of electrodes also varied from patient to patient (*cf.* Table I). The data was recorded from subdural or intracerebral depth electrodes at *f_s_* = 500Hz over the course of several days per patient, and seizures were manually annotated by electroencephalographers, inspecting both raw iEEG and video recordings of the patient. The iEEG signal was notch-filtered at 60Hz and related harmonics to remove powerline trends, and then filtered with an order-3 low-pass filter at 100Hz to remove any high-frequency noise. Then, each electrode channel was independently re-referenced to have zero mean and rescaled to have unit variance.

**TABLE I:**
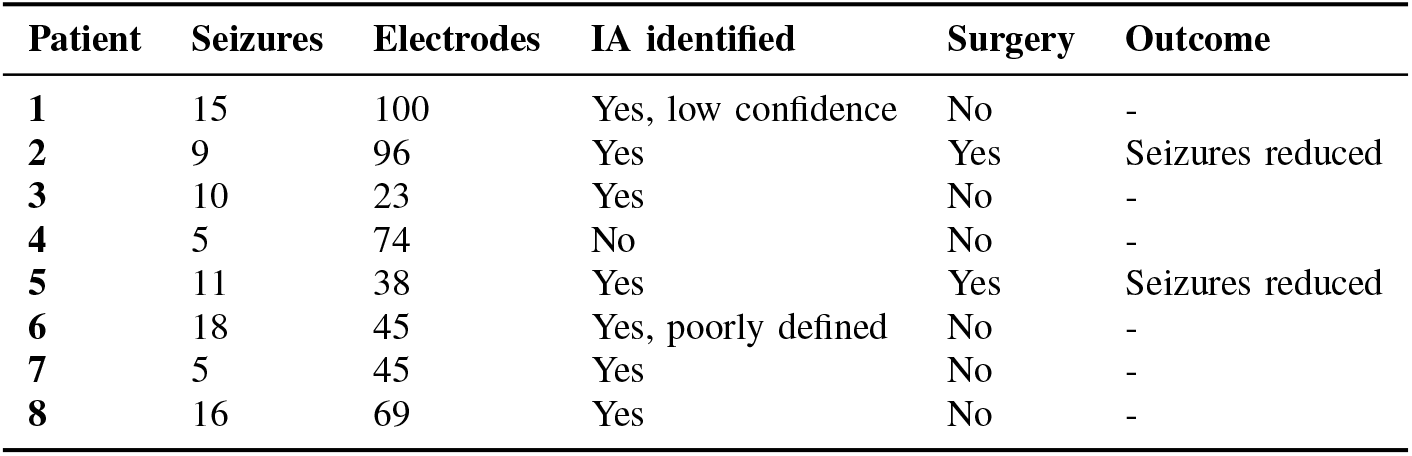
Summary of the patients considered for this study. The columns indicate (left-to-right): the number of recorded seizures, the number of implanted electrodes, the presence of ictal activity (IA) marked by electroencephalographers on one or more channels, whether the patient had surgery, and the outcome of the surgery.

| Patient | Seizures | Electrodes | IA identified | Surgery | Outcome |
| --- | --- | --- | --- | --- | --- |
| 1 | 15 | 100 | Yes, low confidence | No | - |
| 2 | 9 | 96 | Yes | Yes | Seizures reduced |
| 3 | 10 | 23 | Yes | No | - |
| 4 | 5 | 74 | No | No | - |
| 5 | 11 | 38 | Yes | Yes | Seizures reduced |
| 6 | 18 | 45 | Yes, poorly defined | No | - |
| 7 | 5 | 45 | Yes | No | - |
| 8 | 16 | 69 | Yes | No | - |

Before pre-processing, we visually inspected the raw data of each patient and each seizure to assess the presence of bad channels: we considered symmetric windows around each labelled seizure onset and we removed from the data any channels that exhibited abnormal (*i.e.*, either flat or excessive) activity in at least one seizure.

### B. Per-patient analysis of the SOZ

This section reports the available clinical data for the patients considered in our study. For all patients, both the seizure onset time instants and the SOZ annotations were provided by electroencephalographers.

Patient 1 demonstrated ictal activity in both the left and right posterior interhemispheric regions (Figure 3a), with interictal epileptiform discharges recorded independently from the left anterior frontal and right middle frontal lobes. The patient did not undergo resective surgery due to a low confidence in the identification of the SOZ. Patient 2 showed clear seizures originating in the right posterior insular region (Figure 3b). The patient underwent laser interstitial thermal therapy targeting a focal cortical dysplasia in the area. The patient continued to have some post-operative seizures, although these were reduced in frequency and intensity, indicating that the SOZ was identified correctly. Patient 3 had seizure onsets recorded independently from both temporal lobes and thus was not a candidate for surgery. Patient 4 had no clear ictal activity identified by electroencephalographers in the iEEG recordings and was thus not a candidate for surgery, the SOZ evidently not captured by the intracranial electrode placements. Patient 5 demonstrated ictal activity in the left hippocampal body, and underwent a left anterior temporal resection. The patient continued to have seizures after the surgery, but of reduced frequency and intensity, indicating a successful localisation of the SOZ. Patient 6 had multiple seizures recorded with poorly defined, inconsistent ictal onsets over the temporoparietal sensory cortex and was deemed not a candidate for surgical resection due to uncertainty on the SOZ. Patient 7 had seizures recorded in the left hemisphere, with onsets involving a broad region of the temporal lobe neocortex. The patient was not subject to resection due to the epileptogenic zone being too large, and near eloquent language cortex. Patient 8 exhibited abnormal activity in the left amygdala and hippocampus. The patient had already undergone contralateral right anterior temporal resective surgery years prior to the collection of the iEEG data and was not a candidate for further resections.

**Fig. 3:**
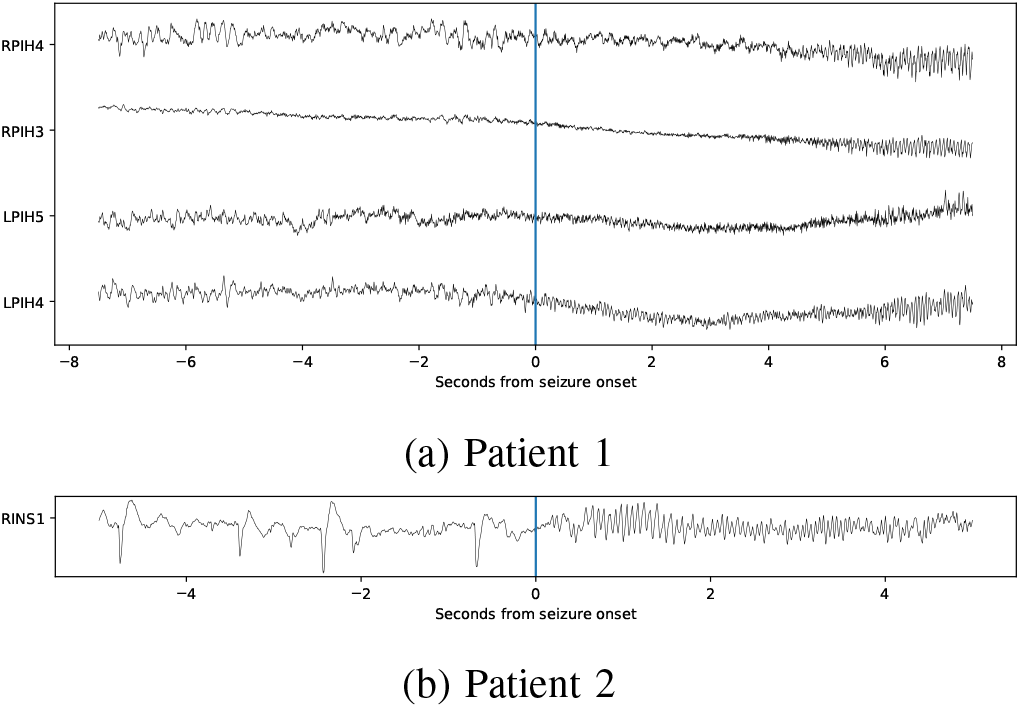
Examples of raw iEEG traces for patients 1 and 2. The two plots show the activity of electrodes that were identified as SOZs by electroencephalographers. The vertical line marks the seizure onset, as reported in the patients’ clinical records.

Table I summarises the relevant details of the eight patients. In particular, six patients had clinically identified, well-defined information regarding the SOZ, whereas in two patients the SOZ could not be clearly identified in the iEEG data by electroencephalographers. Despite not having ground truth information related to the SOZ for these two patients, we still included them as part of our study to analyse the behaviour of our algorithm in such cases of high uncertainty. The question that we aim to answer with this analysis is: what does the GNN see when professional electroencephalographers are uncertain about the SOZ? A strong attention score in such cases would raise concerns about the soundness of our method. Instead, we observe in the following Section that the GNN shows uncertainty in those cases where professionals are also uncertain. This is a valuable result that, in our opinion, strengthens the contributions of the paper.

### C. Results on seizure detection and localisation

Table II reports the Area Under the Receiver Operating Characteristic Curve (ROC-AUC) and the Area Under the Precision-Recall Curve (PR-AUC) obtained by the GNN on the seizure detection task. We report the results obtained using both FC metrics (correlation and PLV) to generate the FNs. We also report the detection performance of a baseline convolutional neural network for time series classification (details in the Appendix). We repeat each experiment five times and, where appropriate, report the average and standard deviation of the results.

**TABLE II:** Average ROC-AUC score and average PR-AUC score for seizure detection on unseen test data. These scores represent the model’s ability to correctly classify the FNs as interictal or ictal. The last row reports the average score over all patients. The highest ROC-AUC and PR-AUC scores are reported in bold for each patient. We report the average and standard deviation over all test seizures and all repetitions.

| Patient | Baseline |  | GNN Corr. |  | GNN PLV |  |
| --- | --- | --- | --- | --- | --- | --- |
|  | ROC | PR | ROC | PR | ROC | PR |
| 1 | 62.54 ± 22.5 | 70.06 ± 17.8 | 68.63 ± 11.43 | 75.20 ± 10.30 | **75.68** ± 23.3 | **77.51** ± 20.1 |
| 2 | 80.19 ± 15.5 | 85.96 ± 10.6 | **86.87** ± 9.07 | **89.04** ± 9.35 | 65.36 ± 20.1 | 72.91 ± 14.8 |
| 3 | 82.32 ± 14.19 | 87.25 ± 9.24 | **93.35** ± 3.12 | **94.34** ± 2.72 | 71.50 ± 14.8 | 71.02 ± 16.3 |
| 4 | 67.81 ± 8.75 | 69.83 ± 13.12 | **60.40** ± 14.41 | **61.11** ± 14.82 | 53.83 ± 6.6 | 51.67 ± 6.4 |
| 5 | 76.18 ± 15.41 | 80.42 ± 14.26 | **77.04** ± 11.98 | **76.39** ± 13.03 | 71.46 ± 12.1 | 71.45 ± 12.9 |
| 6 | **76.32** ± 17.2 | **80.94** ± 13.5 | 73.72 ± 17.14 | 76.02 ± 14.53 | 63.81 ± 17.2 | 71.06 ± 12.4 |
| 7 | 76.46 ± 11.24 | 81.22 ± 7.65 | **85.52** ± 10.95 | **85.92** ± 13.65 | 69.32 ± 2.6 | 65.55 ± 1.8 |
| 8 | 85.60 ± 14.6 | 89.29 ± 10.7 | **90.97** ± 5.51 | **91.89** ± 3.49 | 77.69 ± 11.5 | 78.32 ± 11.3 |
| Avg. | 75.93 ± 7.06 | 80.62 ± 6.86 | **79.56** ± 10.82 | **81.24** ± 10.37 | 68.58 ± 7.08 | 69.94 ± 7.86 |

The GNN achieved an average ROC-AUC score of 79.56 and an average PR-AUC of 81.24 (the average is computed over all patients) when using correlation as FC metric. These results are aligned with the performance of the baseline, which our method slightly outperformed on average, and indicate that 1) our choice of architecture was reasonable and 2) using graph-structured data is an interesting direction for future research on efficient seizure detection. We also recall that the detection task is only meant to provide a weak supervision for the more interesting challenge of localisation, and that better detection results could be achieved by increasing the capacity of the GNN or collecting more training data.

Tables III and IV report the performance of the model on the patients with a known SOZ, respectively using correlation and PLV to generate FNs. In particular, we report three main performance measures:

a. the average precision at *K* (AP@*K*) (Sanderson et al., 2010) obtained by the GNN when computing an average ranking of the electrodes. Each electrode is re-ranked by considering five models trained on the same data and taking the average score assigned to each electrode over all models and all seizures. This measure quantifies the GNN’s ability to correctly identify the SOZ for a patient in general, which is the most clinically relevant scenario.
b. The mean AP@*K* (MAP@*K*) obtained by the GNN on different individual seizures. In this case, the ranking for each seizure is compared to the ground truth independently of the others (*i.e.*, without averaging the scores), and the scores are averaged *a posteriori* (also considering five repetitions of the experiments). This measure quantifies the GNN’s ability to correctly identify target electrodes in a given seizure.
c. The MAP@*K* obtained by the GNN on different individual seizures, but considering groups of electrodes belonging to the same strip (implying spatial locality of the electrodes). This allows us to evaluate the performance of the model at a coarser scale.

From the results we see that, while correlation was a clearly better metric for the task of seizure detection, the localisation performance can vary depending on the particular FC metric used. In particular, the localisation for patients 1 and 5 was better when using correlation networks, but PLV yielded better results for patients 3, 7, and 8.

In general, however, we note that the (M)AP@5 score is positive for both FC metrics, for all performance measures and all patients, meaning that at least one SOZ-associated electrode was ranked in the top five every time. We also note that the GNN achieves a perfect AP@2 score (average rankings) in six out of eight cases when using PLV, indicating a high chance of localising at least two relevant electrodes per patient.

Remarkably, we see that these results were obtained even when considering small datasets, *e.g.*, down to only five seizures for patient 7 (*cf.* Table I). While this result is encouraging and highlights the sample efficiency of our approach, we stress that a higher amount of training data can only improve the detection and, likely, localisation performance of our method, as well as giving a higher statistical certainty about the results.

### D. Comparison with clinical information

Figure 5 shows a graphical visualisation of the scores and rankings used to compute the values in Tables III and IV. The figure summarises our results and provides an overview of the importance scores, their variability across different models and seizures, and their agreement with the ground truth. For every electrode, we report the average score and its standard deviation over all test seizures and all repetitions.

The results for patient 5 can be considered a complete success, with the highest AP@*K* scores among all patients and very little uncertainty in the ranking by the GNN. Crucially, the successful post-operative outcome confirms that the localisation of the SOZ for this patient was accurate and points to a strong localisation ability of the GNN. For patient 2, ictal activity was evident and well-localised on a specific depth electrode placed in the right insular complex (RINS1). The clinical localisation of the SOZ was therefore likely accurate, even if the outcome of the surgery was not completely successful. More importantly, we notice that the GNN was strongly aligned with the human analysis given the same information, and similarly focused on the same electrode (which is ranked first using either of the FC metrics). Our methodology also confirms the conclusions reached by electroencephalographers for patients 3, 7 and 8, although further studies would be required to give a more precise interpretation of the results (including, possibly, the outcome of future surgeries). The results for patient 8 are particularly uncertain, despite the GNN achieving a good detection accuracy (*cf.* Table II). In general, however, the rankings provided by the GNN show a high agreement with the medical assessment in those cases where the SOZ was successfully identified.

For patients with no known SOZ (4, 6) the GNN has a low detection performance and the average attention scores assigned by the GNN are uniformly distributed across all electrodes around an average score of 0.5. On the contrary, patients with a known SOZ have a few electrodes that are assigned a majority of the attentional budget. This difference between the two cases is more clearly visualised in Figure 4, which shows the distribution of the scores given to different electrodes at the seizure onset (patient 5 is taken as representative of the case in which the SOZ is known).

**Fig. 4:**
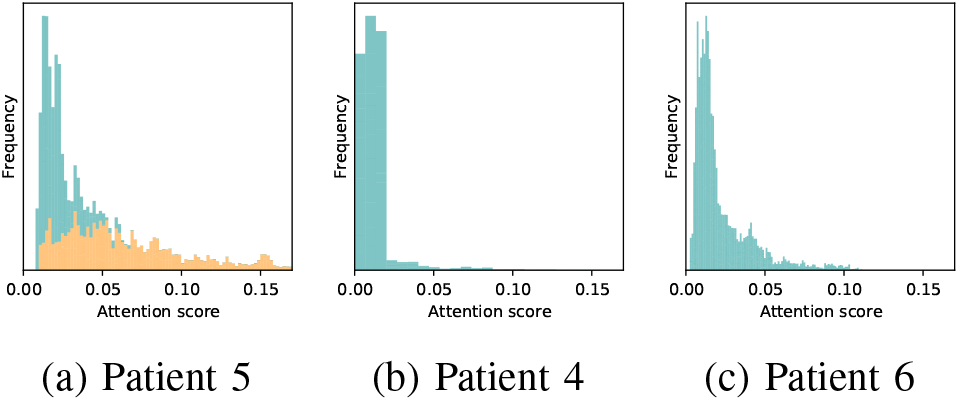
Histograms of the attention scores over a 2-second window starting from a seizure onset. Each bin represents the frequency with which the corresponding attention score is assigned to ten randomly-selected electrodes. Figure **(a)** shows a patient with a known SOZ, while Figures **(b)** and **(c)** show patients without a known SOZ. For Figure **(a)**, the contribution to each bin of those electrodes that are part of the SOZ ground truth are highlighted in orange. Note how the score distribution for SOZ-associated electrodes is spread out towards higher values, while for patients with no known SOZ the scores are similar for all channels.

**Fig. 5:**
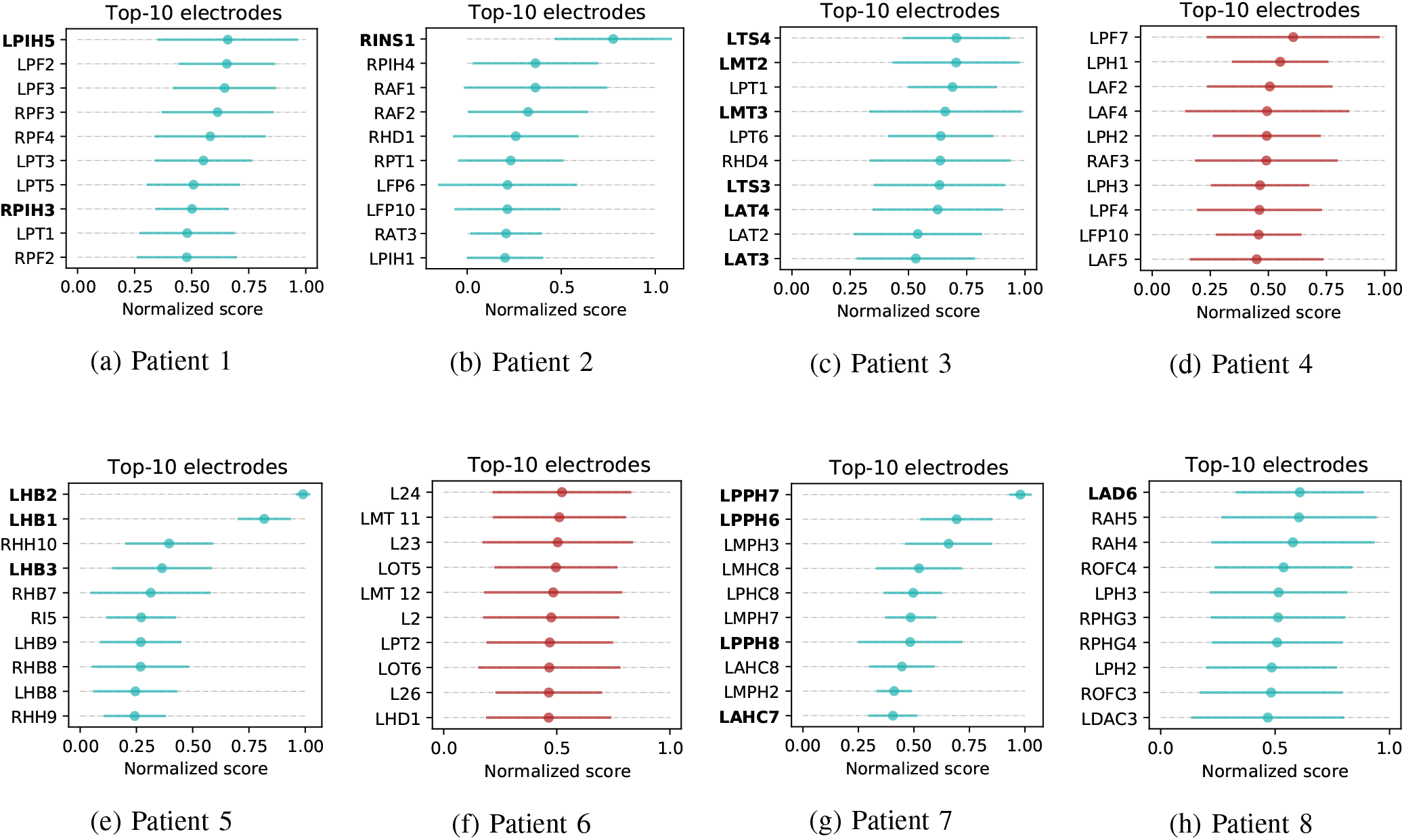
Top ten electrodes when considering the averaged rankings. We report the ranking obtained with the best-performing FC metric for each patient, according to the AP@10 score for average rankings reported in Tables III and IV. The two plots in red indicate those patients for which the SOZ was not identified clinically. Bold labels indicate that the corresponding electrode was marked as a potential SOZ by electroencephalographers. For every electrode, we report the average score and its standard deviation over all test seizures and all repetitions. We refer the reader to the appendix for an extended version of this figure.

For patient 1, the GNN did not identify any particularly important regions despite there being some clinical evidence of ictal activity in the posterior interhemispheric region. Two posterior inter-hemispheric electrodes are indeed ranked in the top ten (averaged rankings) by the GNN when using correlation FNs, although with very high uncertainty. We note, however, that the uncertainty showed by the GNN was also reflected clinically in the electroencephalographers’ interpretations and in the final decision to not operate on this patient.

Our analysis for patients 1, 4, and 6 shows that the uncertainty of the GNN correlates with uncertainty or inability on the part of electroencephalographers to identify the SOZ in iEEG, and can still be useful to support their decision making (*e.g.*, deciding to not operate a patient can be just as valuable as a successful localisation).

## IV. Discussion

Our work introduces a methodology for automated seizure localisation using graph-based machine learning. Our approach does not require any manual annotation of the SOZ in order to work, making it cheaper to train and easier to scale to a larger number of patients. Our method is also data-efficient: we were able to provide a good – and clinically verified – localisation using as little as five annotated seizures per patient.

The goal of the proposed approach is to provide a support tool for clinicians to allocate precious resources in the analysis of iEEG data, and to improve the efficiency of the decision-making process. Crucially, in this regard, we note that our algorithm is conservative in scoring potential SOZ candidates. When the SOZ was not identifiable by electroencephalographers, the GNN also showed uncertainty in the scoring (rather than making high-confidence predictions). Contrarily, a high importance score consistently correlated with clinically-identified SOZs. With this premise, we believe that our approach could have practical value if deployed to epilepsy monitoring units to provide real-time analysis of iEEG recordings.

### A. Future work

There are several directions for future research that could stem from this work. First, we note that by 1) increasing the capacity of the network (in terms of parameters and depth), 2) performing a patient-specific hyperparameter search, and 3) having more seizures on which to train the model, it is likely that both the detection and localisation performance would significantly improve. Also, a possible extension of the proposed methodology could be to explicitly introduce a supervised objective to train the attentional readout using the available information on the SOZ. This would require a per-seizure annotation of every electrode (or, even better, an annotation over time), but could lead to a more accurate localisation. An interesting application of this methodology could also be to provide a patient-agnostic localisation, by training the GNN concurrently on seizures of different patients.

Our current study focuses on eight patients, six of which have an identifiable SOZ. For two of these, we have post-surgical confirmation of the SOZ. Our results are encouraging, but studies on larger sample (with possibly longer-term clinical information on the patients) is required before recommending our approach for clinical practice.

Future work could also explore more in-depth the use of different or combined FC metrics and their impact on the detection and localisation performance. For example, we have observed that using correlation leads to a better detection performance, while we had better localisation results when using PLV. Correlation is the simplest measure for non-directed model-based interactions and is more sensitive to outliers. This sensitivity may result in less uniform FNs between interictal and ictal periods, making it easier for the GNN to detect seizures. However, we argue that it is also this lack of robustness that makes correlation FNs less suitable for localising the SOZ. On the other hand, frequency-domain functional measures like PLV are better for describing whether different brain areas have a preferred phase difference when engaging in oscillatory coupling Bastos and Schoffelen (2016). Due to the synchronous nature of ictal activity, we can assume that PLV will also better highlight those regions of the brain with consistent coupling during seizures and therefore the GNN will be able to assign a high importance score to those regions. Another reason why PLV could be more suitable for localisation is that the SOZ displays internal synchronous activity but also a desynchronisation from the surrounding areas of the brain, possibly making it easier to identify the SOZ. This is discussed in-depth in a study by Le Van Quyen et al. (2001). A way to identify *a priori* the best FC metric to build FNs for a specific patient could bring significant benefits.

## V. Conclusion

We presented a methodology for unsupervised seizure localisation based on GNNs with an attention mechanism. Our approach takes advantage of a compact representation of brain states as FNs, and uses machine learning methods for graph-structured data to automatically detect those regions of the brain that are important for localising seizure onsets. The main advantage of our approach is that it does not require any *a priori* knowledge of the SOZ. The GNN is not forced to focus on any part of the input FNs but, remarkably, learns to focus on areas of the brain that correlate strongly with the true SOZ. We showed the effectiveness of our method in localising the SOZ on real-world data consisting of iEEG recordings from eight human subjects, using two different FC metrics to compute FNs. Our results show a very high accuracy in localising the SOZ. However, we also observed that the GNN exhibits uncertainty in those cases where human analysis was also uncertain, indicating a reliable and safe behaviour to support decision-making.

We believe that this work represents a step towards AI-aided analysis of iEEG data and could potentially lead to faster and more accurate treatment of epilepsy.

## APPENDIX

## A. Seizure generator from Benjamin et al. (2012)

In this experiment we considered a simple network model of seizure initiation presented by Benjamin et al. (2012), and also used by Lopes et al. (2017, 2020) to study the effect of network structure on the generation of seizures. The model consists of a network of *N* bi-stable oscillators

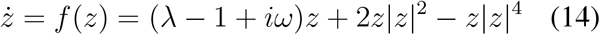

where *z* ∈ ℂ. Equation (14) describes a dynamical system with a stable fixed point at the origin of the complex plane (which we consider as interictal), and an oscillating attractor with frequency *ω* (which we consider as ictal). Parameter *λ* controls the location of the oscillator in phase space. Nodes are interconnected in a graph described by adjacency matrix **A** with a coupling factor *β*, such that the dynamic of a single node reads:

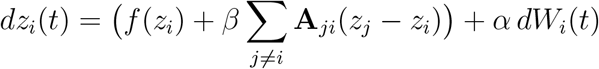

where *W_i_*(*t*) is a stochastic Wiener process rescaled by a factor of *α*.

All nodes in the model are initialised at the fixed point and, due to the presence of noise and the interaction between nodes, eventually switch to the oscillation state. We identify the activity of the whole system as ictal if any of the nodes meets the condition |Re(*z_i_*)| *>* 1, and the SOZ as the first node that escapes the fixed regime.

We consider a complete graph without self-loops to describe the interaction of the nodes. The configuration of the parameters is summarised in Table V. The hyperparameters used for creating the FNs and training the GNN are the same ones that we used for the real iEEG data, and we only report results obtained using PLV as FC metric.

The GNN achieves an almost perfect detection score with a ROC-AUC of 99.61 ± 0.0 and a PR-AUC of 99.69 ± 0.0 (averaged over five runs, evaluated on hold-out test data). Figure 6 compares the generated node activity with the attention scores assigned by the GNN over time. The SOZ channel (green) is assigned the highest attention since the beginning of the seizure until all nodes are simultaneously oscillating, at which point the attention scores converge to be evenly distributed. A similar even distribution is observed in the interictal state, indicating that the network has correctly learned to identify the SOZ electrode without defaulting to assign a high score to just one electrode. This behaviour is confirmed by the spikes in attention assigned to channels 0 and 1, which happen as soon as the node dynamics escape the fixed-point attractor.

**Fig. 6:**
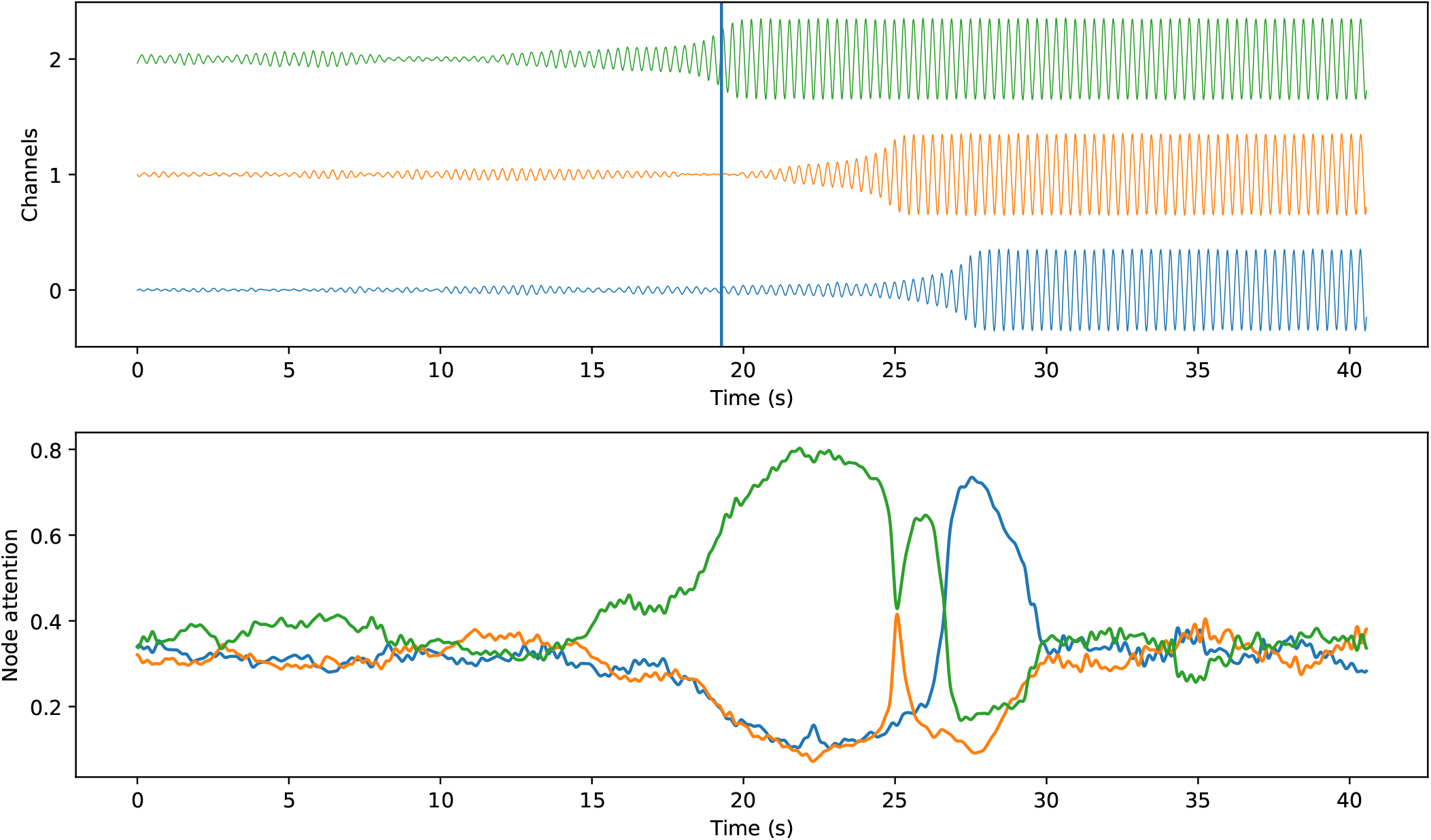
Top: a clip showing the generated activity of a 3-node simulator, compared to the attention coefficient assigned by the GNN at each node over time. Colours indicate the same node in both plots.

## B. The Virtual Brain Simulator

In this experiment we use The Virtual Brain simulator (TVB) (Sanz Leon et al., 2013) to model a patient with temporal lobe epilepsy.

We follow the same approach described in TVB’s documentation to configure the simulator.^1^ We assign the Epileptor neural mass model (Jirsa et al., 2014) to all the controllable brain regions of TVB. We set the epileptogenicity of the right limbic areas (rHC, rPHC and rAMYG) to −1.6, the superior temporal cortex (rTCI) and the ventral temporal cortex (rTCV) to −1.8, while for all other areas to −2.2. The remaining parameters are kept as default. The hyperparameters used for creating the FNs and training the GNN are the same ones that we used for the real iEEG data.

We select a subset of 34 sEEG virtual sensors among the ones provided for the default subject of TVB. Of this subset, electrode 33 shows strong epileptogenic activity, while electrodes 18, 19, and 20 show mild activity. We generate clips of roughly 1 minute at 20Hz so that there is a simulated onset in the middle of each clip. An example of a generated clip is shown in Figure 7.

**Fig. 7:**
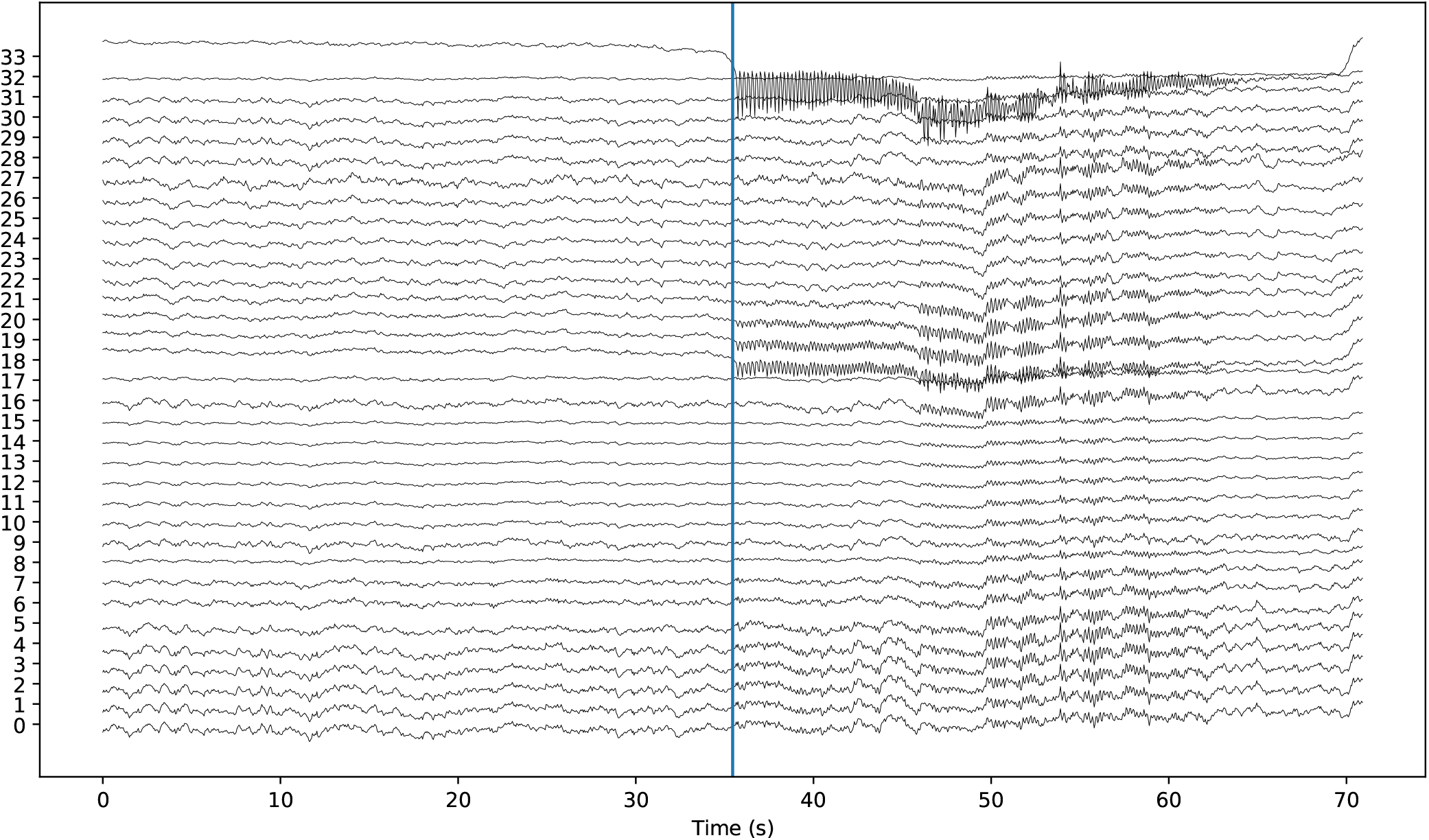
A virtual seizure generated with TVB. The vertical line denotes the annotated seizure onset in time.

The GNN achieved an average detection ROC-AUC of 98.87 ± 0.18 and an average PR-AUC of 99.18 ± 0.07 (averaged over five runs, evaluated on hold-out test data). The electrode with a strong ictal activity is consistently assigned a maximum score of 1 by all models and electrode 19 is also ranked in the top-5 electrodes (see Figure 8).

**Fig. 8:**
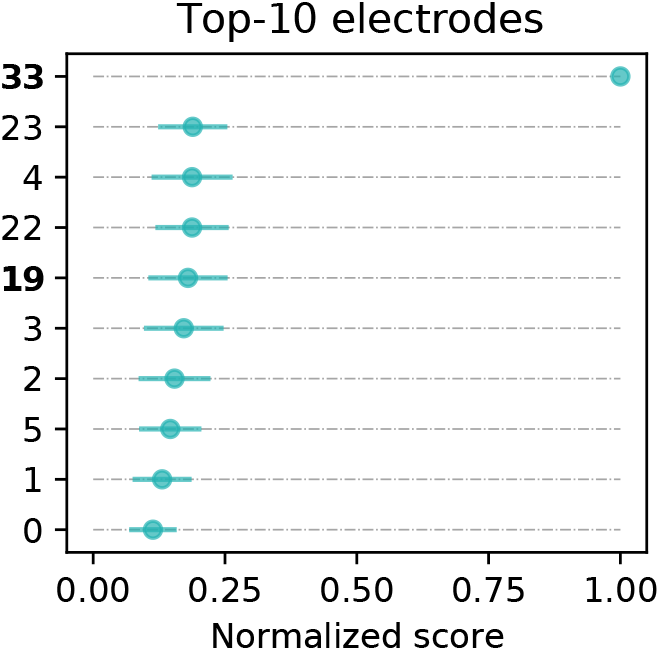
Top-10 electrodes with averaged rankings. Bold labels indicate that the corresponding electrode showed ictal activity. As desired, electrode 33 shows strong epileptogenic activity.

## C. GNN training details

We consider each patient separately and train a GNN from scratch to build patient-specific models. The GNN architecture is the one given in Equation (11). The ECC layer has 32 output units with ReLU activation and a kernel-generating network *f* (·) consisting of a two-layer MLP with 32 hidden units and ReLU activation. All parameters of the layer are regularised with an *L*_2_ penalty with a factor of 10^*−*5^.

The MLP classifier following the ATTN-RO readout has 2 layers, with the hidden one having 32 units and ReLU activation and with 25% dropout in-between. Both layers are regularised with an *L*_2_ penalty with factor 10^*−*5^.

The model is trained using Adam, with a learning rate of 10^*−*3^ and a batch size of 32 graphs. The model is trained to convergence with 10 epochs of patience, using the data from [0.1·*n*] seizures selected randomly (*n* being the overall number of seizures) for early stopping. We then test the model on a held-out set of [0.1·*n*] seizures. The remaining seizures are used for training. All experiments are repeated 5 times using different random data splits.

## D. Baseline training details

The baseline is a simple 1D convolutional neural network (CNN) based on the architecture described by Wang et al. (2017). The CNN operates directly on iEEG time series and hence does not take into account any graph-based representation for the data. Similarly to how we create the input-output pairs for the GNN, here we consider windows of size *T* taken at a stride of *k/f_s_* for the interictal class and stride 1*/f_s_* for the ictal class, and we associate to each window a class label corresponding to the majority class of *y*(*t*) in the corresponding window.

In particular, we shrink the model to make it comparable in terms of number of parameters and depth to the GNN one, and also to prevent over-fitting (which we experimentally encountered as a significant problem with the model). We consider a single convolutional layer with a kernel of size 3, 8 output channels, and ReLU activations, followed by a global average pooling and a single-layer MLP to output the classification decision. We train the model using Adam with learning rate 0.001, batch size of 32 and early stopping with a patience of 5 epochs.

## E. Additional results

## Detection

A notable behaviour of the GNN can be observed from Figure 9, which shows the output of the GNN (*i.e.*, the detection score outputted by the model) on a symmetrical window around the onset, for randomly sampled seizures of the six patients with a known SOZ. We empirically observed that the model is robust to the onset labelling provided by electroencephalographers. Notably, by analysing the prediction of the GNN in the time instants prior to the seizure onset, we can see that the confidence with which the GNN detects a seizure starts to gradually increase towards the seizure onset, but does not always peak at the onset time marked by electroencephalographers. This suggests that the GNN is learning to detect the anomalous brain activity rather than overfitting to the known onset labels.

**Fig. 9:**
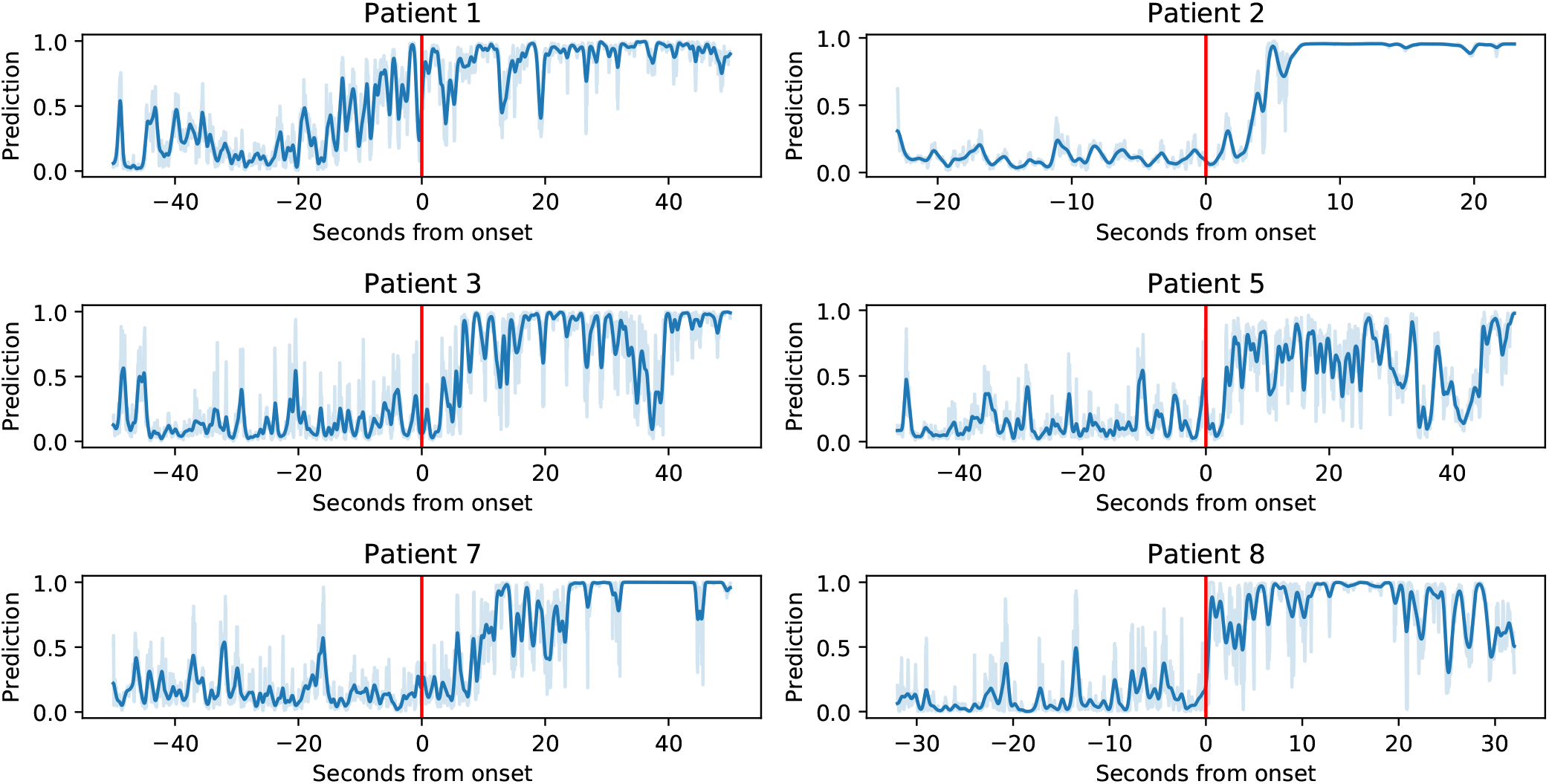
Example of the detection score outputted by the GNN, for all patients with a known SOZ. We show a window of 50 seconds around the marked onset for random test seizures. The darker line is a smoothed trendline of the true prediction, shown in lighter colour.

## Localisation

We show in Figures 11 and 12 the top 10 electrodes by AP@10 score for all patients, respectively when using correlation and PLV as FC metrics.

## Threshold

To demonstrate that our approach is robust to the choice of sparsification threshold for the FNs, as argued in Section II-A, We report in Figure 10 the average localisation performance over all patients and all metrics for different thresholds (that is, we average all the values reported in Table III after re-computing the tables with different sparsification thresholds). While this is a coarse-grained analysis, it shows that there are no significant differences in the downstream performance for thresholds up to 0.7, with two-sided t-tests over all pairs yielding p-value *p* ≫ 0.05 up until threshold 0.7. Above this value, we see a significant performance degradation.

**Fig. 10:**
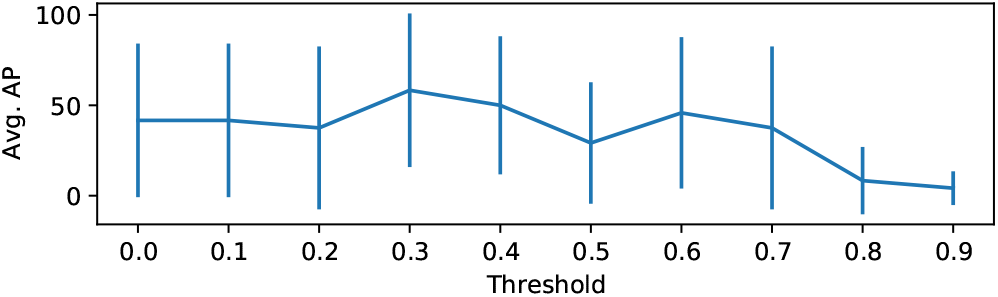
Average detection and localisation performance as a function of the sparsification threshold. We report the average over all metrics and all patients, as reported in Tables II and III of the manuscript.

**Fig. 11:**
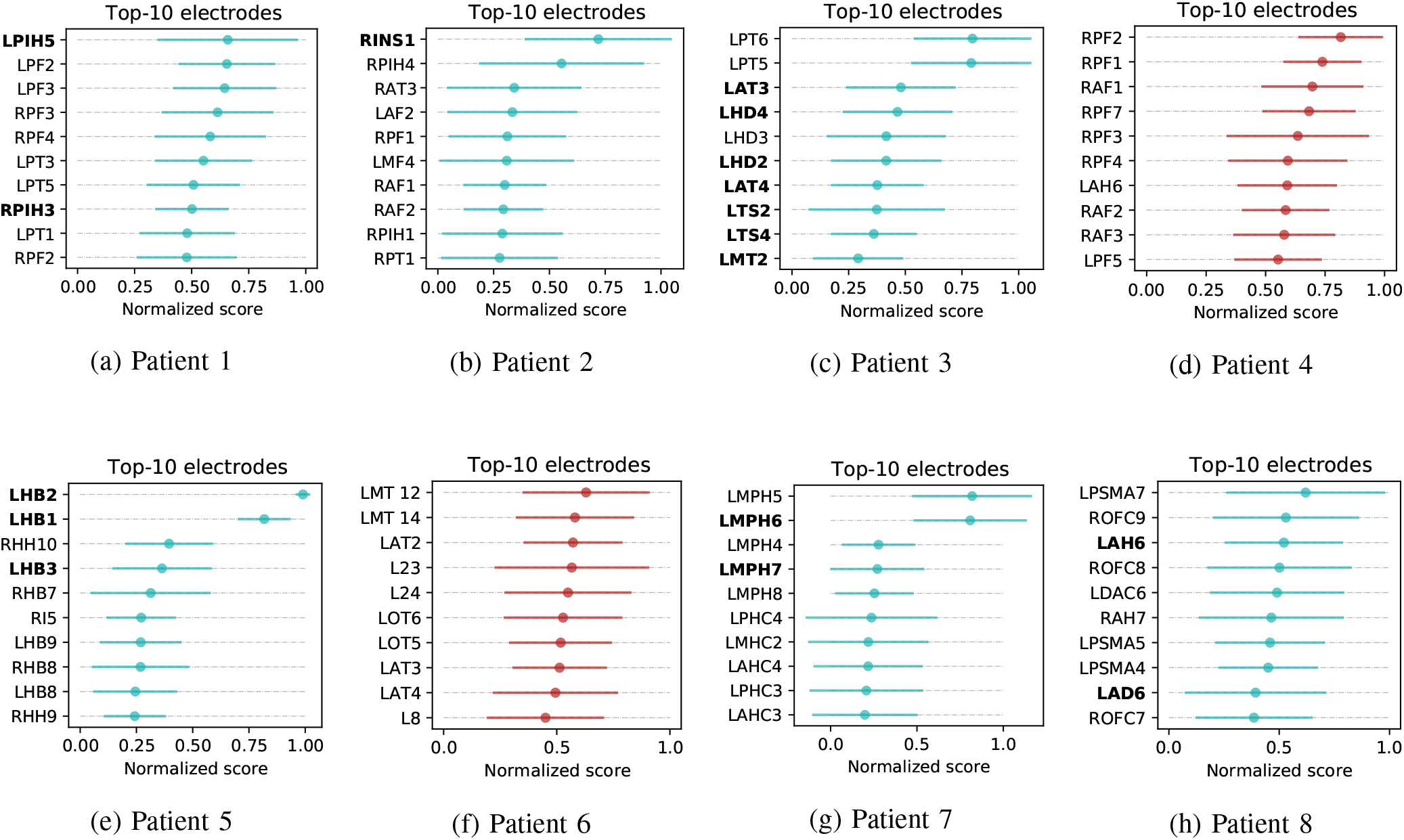
Top ten electrodes by AP@10 score for the average rankings, using correlation as FC measure. The two plots in red indicate those patients for which the SOZ was not identified clinically. Bold labels indicate that the corresponding electrode was marked as a potential SOZ by electroencephalographers.

**Fig. 12:**
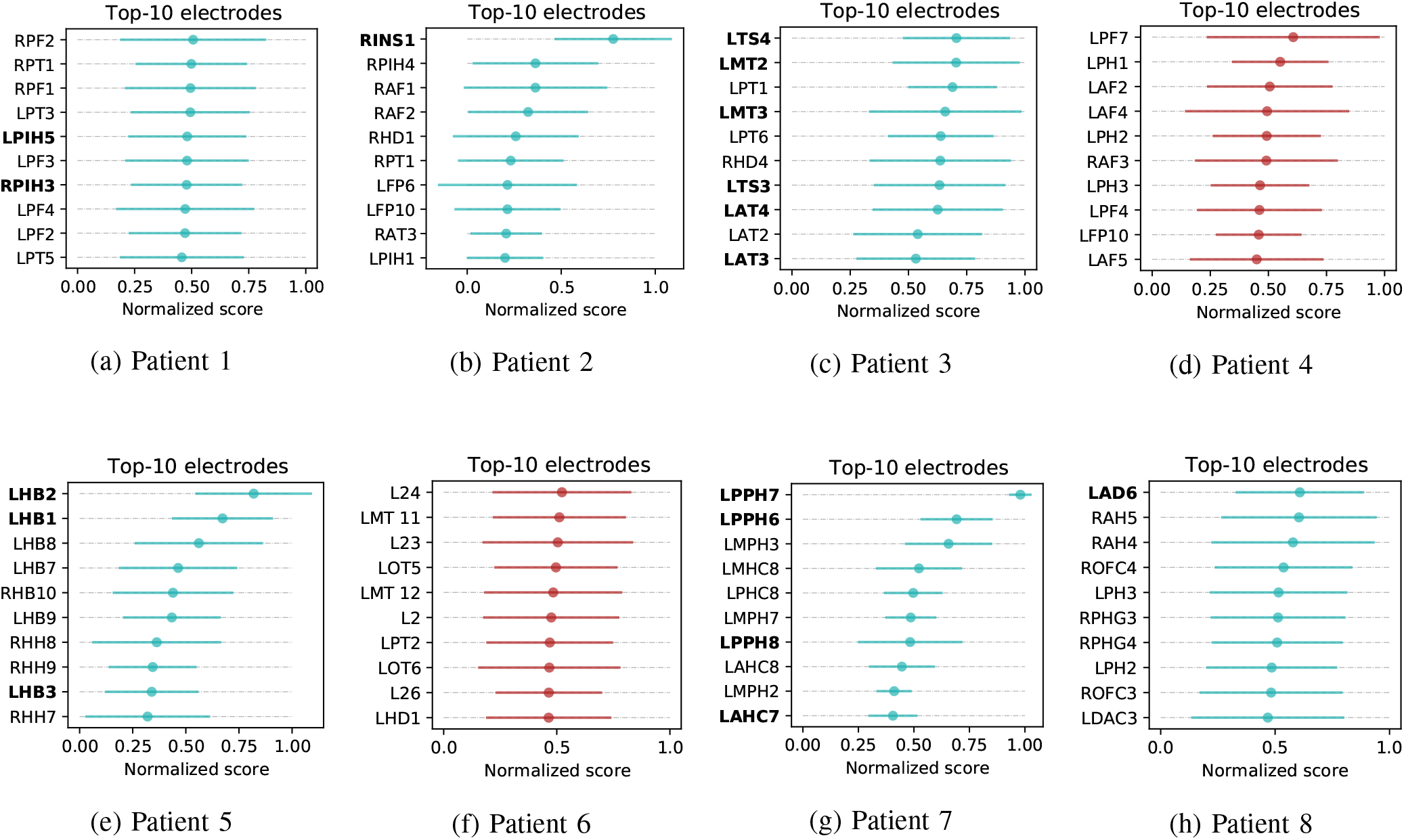
Top ten electrodes by AP@10 score for the average rankings, using PLV as FC measure. The two plots in red indicate those patients for which the SOZ was not identified clinically. Bold labels indicate that the corresponding electrode was marked as a potential SOZ by electroencephalographers.

**TABLE III:**
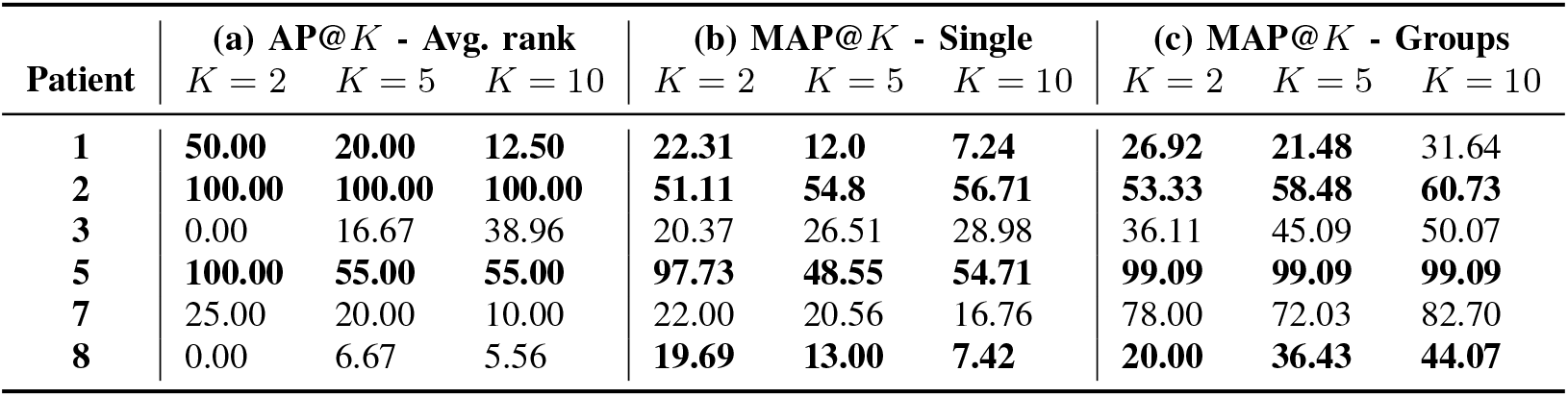
Localisation performance for patients with a known SOZ, when using Pearson’s correlation as FC metric. We report: **(a)** the average precision at *K* for averaged rankings, which evaluates the localisation for the patient overall; **(b)** the mean average precision at *K* for single rankings, which evaluates the localisation for a given seizure; **(c)** the mean average precision at *K* for single rankings and groups of electrodes, which is equivalent to (b) but at a coarser scale. We report scores for *K* = 2, 5, 10. Bold indicates that the results are better than the ones obtained with PLV as FC metric (*cf.* Table IV).

| Patient | (a) AP@$K$ - Avg. rank | | | (b) MAP@$K$ - Single | | | (c) MAP@$K$ - Groups | | |
| --- | --- | --- | --- | --- | --- | --- | --- | --- | --- |
| | $K = 2$ | $K = 5$ | $K = 10$ | $K = 2$ | $K = 5$ | $K = 10$ | $K = 2$ | $K = 5$ | $K = 10$ |
| 1 | **50.00** | **20.00** | **12.50** | **22.31** | **12.0** | **7.24** | **26.92** | **21.48** | 31.64 |
| 2 | **100.00** | **100.00** | **100.00** | **51.11** | **54.8** | **56.71** | **53.33** | **58.48** | **60.73** |
| 3 | 0.00 | 16.67 | 38.96 | 20.37 | 26.51 | 28.98 | 36.11 | 45.09 | 50.07 |
| 5 | **100.00** | **55.00** | **55.00** | **97.73** | **48.55** | **54.71** | **99.09** | **99.09** | **99.09** |
| 7 | 25.00 | 20.00 | 10.00 | 22.00 | 20.56 | 16.76 | 78.00 | 72.03 | 82.70 |
| 8 | 0.00 | 6.67 | 5.56 | **19.69** | **13.00** | **7.42** | **20.00** | **36.43** | **44.07** |

**TABLE IV:** Localisation performance for patients with a known SOZ, when using PLV as FC metric. Bold indicates that the results are better than the ones obtained with correlation as FC metric (*cf.* Table III).

| Patient | (a) AP@$K$ - Avg. rank | | | (b) MAP@$K$ - Single | | | (c) MAP@$K$ - Groups | | |
| --- | --- | --- | --- | --- | --- | --- | --- | --- | --- |
| | $K = 2$ | $K = 5$ | $K = 10$ | $K = 2$ | $K = 5$ | $K = 10$ | $K = 2$ | $K = 5$ | $K = 10$ |
| 1 | 0.00 | 5.00 | 5.83 | 8.67 | 5.97 | 4.67 | 16.67 | 18.82 | **31.78** |
| 2 | **100.00** | **100.00** | **100.00** | 50.00 | 54.33 | 56.65 | 50.00 | 58.04 | 60.21 |
| 3 | **100.00** | **55.00** | **45.46** | **60.00** | **40.82** | **32.58** | **66.88** | **45.16** | **51.36** |
| 5 | **100.00** | 40.00 | 48.57 | 66.82 | 38.28 | 45.27 | 91.82 | 93.48 | 93.48 |
| 7 | **100.00** | **40.00** | **35.71** | **70.00** | **43.84** | **30.45** | **82.00** | **74.22** | **84.22** |
| 8 | **50.00** | **20.00** | **10.00** | 15.62 | 9.15 | 6.43 | 16.56 | 20.07 | 32.30 |

**TABLE V:** Configuration used for the simulator by Benjamin et al. (2012).

| Parameter | Value |
| --- | --- |
| $N$ | 3 |
| $\omega$ | 20 |
| $\lambda$ | 0.5 |
| $\beta$ | 0.1 |
| $\alpha$ | 0.05 |

While this is a coarse-grained analysis, it indicates that the most meaningful edges to perform seizure localisation are those that indicate a strong functional connectivity, with values higher than 0.7. At the same time, a higher sparsification threshold can improve the computational cost of the GNN, which is linear in the number of edges. However, it is beyond the scope of this work to provide a biological interpretation of this threshold and we do not make claims regarding the generality of this threshold. Our general recommendation, if computational cost is not a priority, is to keep the threshold conservatively low so as to not remove potentially informative edges from the FNs. The value of 0.1 that we use in our experiments appears to be a reasonable choice, although we leave further exploration of this matter as future work.

## Footnotes

1 https://github.com/the-virtual-brain/tvb-root/blob/master/tvb_documentation

## References

C. E. Stafstrom and L. Carmant, “Seizures and epilepsy: An overview for neuroscientists,” Cold Spring Harbor Perspectives in Medicine, vol. 5, no. 6, p. a022426, 2015.

P. Kwan and M. J. Brodie, “Early identification of refractory epilepsy,” New England Journal of Medicine, vol. 342, no. 5, pp. 314–319, 2000.

S. P. Burns, S. Santaniello, R. B. Yaffe, C. C. Jouny, N. E. Crone, G. K. Bergey, W. S. Anderson, and S. V. Sarma, “Network dynamics of the brain and influence of the epileptic seizure onset zone,” Proceedings of the National Academy of Sciences, vol. 111, no. 49, pp. 321–330, 2014.

P. Van Mierlo, M. Papadopoulou, E. Carrette, P. Boon, S. Vandenberghe, K. Vonck, and D. Marinazzo, “Functional brain connectivity from eeg in epilepsy: Seizure prediction and epileptogenic focus localization,” Progress in Neurobiology, vol. 121, pp. 19–35, 2014.

P. L. Nunez, R. Srinivasan et al., Electric fields of the brain: the neurophysics of EEG. Oxford University Press, USA, 2006.

A. K. Shah and S. Mittal, “Invasive electroencephalography monitoring: Indications and presurgical planning,” Annals of Indian Academy of Neurology, vol. 17, no. Suppl 1, p. S89, 2014.

K. Hashiguchi, T. Morioka, F. Yoshida, Y. Miyagi, S. Nagata, A. Sakata, and T. Sasaki, “Correlation between scalp-recorded electroencephalographic and electrocorticographic activities during ictal period,” Seizure, vol. 16, no. 3, pp. 238–247, 2007.

A. M. Bastos and J.-M. Schoffelen, “A tutorial review of functional connectivity analysis methods and their interpretational pitfalls,” Frontiers in Systems Neuroscience, vol. 9, p. 175, 2016.

W. Gersch and G. Goddard, “Epileptic focus location: spectral analysis method,” Science, vol. 169, no. 3946, pp. 701–702, 1970.

M. A. Brazier, “Spread of seizure discharges in epilepsy: anatomical and electrophysiological considerations,” Experimental Neurology, vol. 36, no. 2, pp. 263–272, 1972.

A. N. Khambhati, K. A. Davis, B. S. Oommen, S. H. Chen, T. H. Lucas, B. Litt, and D. S. Bassett, “Dynamic network drivers of seizure generation, propagation and termination in human neocortical epilepsy,” PLoS Computational biology, vol. 11, no. 12, p. e1004608, 2015.

A. N. Khambhati, K. A. Davis, T. H. Lucas, B. Litt, and D. S. Bassett, “Virtual cortical resection reveals push-pull network control preceding seizure evolution,” Neuron, vol. 91, no. 5, pp. 1170–1182, 2016.

K. A. Schindler, S. Bialonski, M.-T. Horstmann, C. E. Elger, and K. Lehnertz, “Evolving functional network properties and synchronizability during human epileptic seizures,” Chaos: An Interdisciplinary Journal of Nonlinear Science, vol. 18, no. 3, p. 033119, 2008.

M. A. Lopes, M. P. Richardson, E. Abela, C. Rummel, K. Schindler, M. Goodfellow, and J. R. Terry, “An optimal strategy for epilepsy surgery: Disruption of the rich-club?” PLoS Computational Biology, vol. 13, no. 8, p. e1005637, 2017.

H. W. Lee, J. Arora, X. Papademetris, F. Tokoglu, M. Negishi, D. Scheinost, P. Farooque, H. Blumenfeld, D. D. Spencer, and R. T. Constable, “Altered functional connectivity in seizure onset zones revealed by fmri intrinsic connectivity,” Neurology, vol. 83, no. 24, pp. 2269–2277, 2014.

K. E. Weaver, W. Chaovalitwongse, E. J. Novotny, A. Poliakov, T. J. Grabowski Jr, and J. G. Ojemann, “Local functional connectivity as a presurgical tool for seizure focus identification in non-lesion, focal epilepsy,” Frontiers in Neurology, vol. 4, p. 43, 2013.

W. Staljanssens, G. Strobbe, R. Van Holen, G. Birot, M. Gschwind, M. Seeck, S. Vandenberghe, S. Vulliémoz, and P. van Mierlo, “Seizure onset zone localization from ictal high-density eeg in refractory focal epilepsy,” Brain Topography, vol. 30, no. 2, pp. 257–271, 2017.

I. Covert, B. Krishnan, I. Najm, J. Zhan, M. Shore, J. Hixson, and M. J. Po, “Temporal Graph Convolutional Networks for Automatic Seizure Detection,” arXiv preprint arXiv:1905.01375, 2019.

B. Yu, H. Yin, and Z. Zhu, “Spatio-temporal graph convolutional networks: A deep learning framework for traffic forecasting,” arXiv preprint arXiv:1709.04875, 2017.

S. Gadgil, Q. Zhao, E. Adeli, A. Pfefferbaum, E. V. Sullivan, and K. M. Pohl, “Spatio-temporal graph convolution for functional mri analysis,” arXiv preprint arXiv:2003.10613, 2020.

G. Litjens, T. Kooi, B. E. Bejnordi, A. A. A. Setio, F. Ciompi, M. Ghafoorian, J. A. Van Der Laak, B. Van Ginneken, and C. I. Sánchez, “A survey on deep learning in medical image analysis,” Medical Image Analysis, vol. 42, pp. 60–88, 2017.

P. W. Battaglia, J. B. Hamrick, V. Bapst, A. Sanchez-Gonzalez, V. Zambaldi, M. Malinowski, A. Tacchetti, D. Raposo, A. Santoro, R. Faulkner et al., “Relational inductive biases, deep learning, and graph networks,” arXiv preprint arXiv:1806.01261, 2018.

M. M. Bronstein, J. Bruna, Y. LeCun, A. Szlam, and P. Vandergheynst, “Geometric deep learning: going beyond euclidean data,” IEEE Signal Processing Magazine, vol. 34, no. 4, pp. 18–42, 2017.

O. Benjamin, T. H. Fitzgerald, P. Ashwin, K. Tsaneva-Atanasova, F. Chowdhury, M. P. Richardson, and J. R. Terry, “A phenomenological model of seizure initiation suggests network structure may explain seizure frequency in idiopathic generalised epilepsy,” The Journal of Mathematical Neuroscience, vol. 2, no. 1, pp. 1–30, 2012.

P. Sanz Leon, S. A. Knock, M. M. Woodman, L. Domide, J. Mersmann, A. R. McIntosh, and V. Jirsa, “The virtual brain: a simulator of primate brain network dynamics,” Frontiers in Neuroinformatics, vol. 7, p. 10, 2013.

V. K. Jirsa, W. C. Stacey, P. P. Quilichini, A. I. Ivanov, and C. Bernard, “On the nature of seizure dynamics,” Brain, vol. 137, no. 8, pp. 2210–2230, 2014.

J.-P. Lachaux, E. Rodriguez, J. Martinerie, and F. J. Varela, “Measuring phase synchrony in brain signals,” Human Brain Mapping, vol. 8, no. 4, pp. 194–208, 1999.

M. A. Kramer, U. T. Eden, S. S. Cash, and E. D. Kolaczyk, “Network inference with confidence from multivariate time series,” Physical Review E, vol. 79, no. 6, p. 061916, 2009.

D. Bahdanau, K. Cho, and Y. Bengio, “Neural machine translation by jointly learning to align and translate,” arXiv preprint arXiv:1409.0473, 2014.

A. Vaswani, N. Shazeer, N. Parmar, J. Uszkoreit, L. Jones, A. N. Gomez, L. Kaiser, and I. Polosukhin, “Attention is all you need,” in Advances in neural information processing systems, 2017, pp. 5998–6008.

T. B. Brown, B. Mann, N. Ryder, M. Subbiah, J. Kaplan, P. Dhariwal, A. Neelakantan, P. Shyam, G. Sastry, A. Askell et al., “Language models are few-shot learners,” arXiv preprint arXiv:2005.14165, 2020.

K. Xu, J. Ba, R. Kiros, K. Cho, A. Courville, R. Salakhudinov, R. Zemel, and Y. Bengio, “Show, attend and tell: Neural image caption generation with visual attention,” in International conference on machine learning, 2015, pp. 2048–2057.

P. Velickovic, G. Cucurull, A. Casanova, Romero, P. Lio, and Y. Bengio, “Graph attention networks,” International Conference of Learning Representations (ICLR), 2018.

J. Gilmer, S. S. Schoenholz, P. F. Riley, O. Vinyals, and G. E. Dahl, “Neural message passing for quantum chemistry,” in Proceedings of the 34th International Conference on Machine Learning-Volume 70. JMLR.org, 2017, pp. 1263–1272.

M. Defferrard, X. Bresson, and P. Vandergheynst, “Convolutional neural networks on graphs with fast localized spectral filtering,” in Advances in Neural Information Processing Systems, 2016, pp. 3844–3852.

F. M. Bianchi, D. Grattarola, L. Livi, and C. Alippi, “Graph neural networks with convolutional ARMA filters,” arXiv preprint arXiv:1901.01343, 2019.

M. Simonovsky and N. Komodakis, “Dynamic edgeconditioned filters in convolutional neural networks on graphs,” in Proceedings of the IEEE Conference on Computer Vision and Pattern Recognition, 2017.

M. Schlichtkrull, T. N. Kipf, P. Bloem, R. van den Berg, I. Titov, and M. Welling, “Modeling relational data with graph convolutional networks,” in European Semantic Web Conference. Springer, 2018, pp. 593–607.

M. Sanderson, C. D. Manning, P. Raghavan, and H. Schütze, “Introduction to information retrieval, cambridge university press. 2008. isbn-13 978-0-521-86571-5, xxi+ 482 pages.” Natural Language Engineering, vol. 16, no. 1, pp. 100–103, 2010.

M. Le Van Quyen, J. Martinerie, V. Navarro, M. Baulac, and F. J. Varela, “Characterizing neurodynamic changes before seizures,” Journal of Clinical Neurophysiology, vol. 18, no. 3, pp. 191–208, 2001.

M. A. Lopes, L. Junges, W. Woldman, M. Good-fellow, and J. R. Terry, “The role of excitability and network structure in the emergence of focal and generalized seizures,” Frontiers in Neurology, vol. 11, p. 74, 2020.

Z. Wang, W. Yan, and T. Oates, “Time series classification from scratch with deep neural networks: A strong baseline,” in 2017 International joint conference on neural networks (IJCNN). IEEE, 2017, pp. 1578–1585.

